# *In vivo* tracking of adenoviral-transduced iron oxide-labeled bone marrow-derived dendritic cells using magnetic particle imaging

**DOI:** 10.1101/2023.03.14.532667

**Authors:** Corby Fink, Julia J. Gevaert, John W. Barrett, Jimmy D. Dikeakos, Paula J. Foster, Gregory A. Dekaban

## Abstract

**Background:** Despite widespread study of dendritic cell (DC)-based cancer immunotherapies, the *in vivo* post-injection fate of DC remains largely unknown. Due in part to a lack of quantifiable imaging modalities, this is troubling as the amount of DC migration to secondary lymphoid organs correlates with therapeutic efficacy. Preliminary studies have identified magnetic particle imaging (MPI) as a suitable modality to quantify *in vivo* migration of superparamagnetic iron oxide-(SPIO)-labeled DC. Herein, we describe a lymph node- (LN)-focused MPI scan to quantify DC *in vivo* migration accurately and consistently.

**Methods:** Both adenovirus (Ad)-transduced SPIO^+^ (Ad SPIO^+^) and SPIO^+^ C57BL/6 bone marrow-derived DC were generated and assessed for viability and phenotype using flow cytometry. Ad SPIO^+^ and SPIO^+^ DC were fluorescently-labeled and injected into C57BL/6 mouse hind footpads (n=6). Two days later, *in vivo* DC migration was quantified using whole animal, popliteal LN- (pLN)-focused, and *ex vivo* pLN MPI scans.

**Results:** No significant differences in viability, phenotype and *in vivo* pLN migration were noted for Ad SPIO^+^ and SPIO^+^ DC. Day 2 pLN-focused MPI successfully quantified DC migration in all instances while whole animal MPI only quantified pLN migration in 75% of cases. *Ex vivo* MPI and fluorescence microscopy confirmed MPI signal was pLN-localized and due to originally-injected Ad SPIO^+^ and SPIO^+^ DC.

**Conclusions:** We overcame a reported limitation of MPI by using a pLN-focused MPI scan to quantify pLN-migrated Ad SPIO^+^ and SPIO^+^ DC in 100% of cases. With this improved method, we detected as few as 1000 DC (4.4 ng Fe) *in vivo*. MPI is a suitable pre-clinical imaging modality to assess DC-based cancer immunotherapeutic efficacy.

## INTRODUCTION

For cancer patients who are non-responsive to or metastatically progress after standard of care treatment options, cancer immunotherapy remains an available treatment option [1]. Cancer immunotherapy initiates the enhancement of one’s own immune system to combat cancer. As such, *ex vivo*-generated autologous dendritic cells (DC) can be exploited as professional antigen-presenting cells (APC) to express tumor-associated antigens (TAA) in conjunction with co-stimulatory molecules required to initiate a potent TAA-specific immune response upon injection into the host [2]. Delivery of TAA to DC can occur via different mechanisms, but viral transduction of a TAA-expressing vector remains an efficient delivery system [3]. The immune response elicited by this DC-based cancer vaccine strategy includes the direct activation of TAA-specific CD4^+^ and CD8^+^ T cells in conjunction with indirect activation of innate immune cells, namely natural killer (NK) cells, NK T cells, and macrophages [4]. Despite the potential of DC to orchestrate robust TAA-specific immune responses [5], their treatment effectiveness remains sub-optimal [6] and the *in vivo* fate of therapeutic DC following administration remains largely unknown.

The paucity of knowledge on *in vivo* DC localization is troubling. First, DC migration to secondary lymphoid organs post injection and subsequent presentation of TAA to prime T cells post injection is required and secondly, quantification of this DC migration correlates with therapeutic potency [7, 8]. When considering that a limitation of treatment effectiveness stems from < 5% of injected DC reaching secondary lymphoid organs [6, 9, 10], the necessity for non-invasive quantifiable DC *in vivo* tracking is needed to both assess and improve upon vaccine outcomes.

We and others have employed magnetic resonance imaging (MRI) as a non-invasive imaging modality to track both human and mouse DC *in vivo* in pre-clinical and clinical studies [11–15]. By labeling DC *ex vivo* with superparamagnetic iron oxide nanoparticles (SPIO), SPIO^+^ DC are identifiable *in vivo* as regions of signal hypointensities resulting from the effect of SPIO^+^ DC on surrounding proton relaxation. Although highly sensitive [16], quantification of SPIO^+^ DC with MRI remains semi-quantitative [17] and low in specificity as signal hypointensities inherent in certain tissues cannot be discriminated from SPIO^+^ DC-induced hypointensities [18]. Magnetic particle imaging (MPI) has now emerged as a pre-clinical small animal alternative to MRI cell tracking of SPIO^+^ cells [19]. Unlike MRI, MPI signal is quantifiable as it is linearly proportional to iron mass. Tissue/bone hypointensities that complicate MRI are also non-existent with MPI [20]. Low resolution and biocompatibility of SPIO are noted limitations of MPI [21, 22]; yet hardware upgrades [23] and the development of new MPI-tailored SPIO formulations are ongoing [24]. We previously tracked *in vivo* SPIO^+^ bone marrow-derived DC (BMDC) migration with MPI [14]. However, a major challenge has been the limited dynamic range of MPI when there are two signals of interest in the field of view (FOV) with different iron concentrations. In our pre-clinical model, the high iron concentration from administered BMDC at the injection site shadows the lower iron concentration from BMDC in the draining lymph node (LN), reducing the ability to separate and accurately quantify signals. Due to its broad applicability across a range of disease states and with the first human MPI scanner designs increasingly becoming a reality [25], pre-clinical studies optimizing the tracking of SPIO^+^ BMDC is of timely importance.

Herein, we designed a combined protocol to generate adenoviral-transduced SPIO-labeled BMDC without affecting viability, phenotype or *in vivo* migration to popliteal LN (pLN) detected by MPI. Additionally, by combining the use of a focused FOV centred on pLN with a multichannel joint image reconstruction algorithm, we are the first group to successfully overcome the dynamic range limitation of MPI to permit accurate quantification of pLN-migrated BMDC. MPI is a suitable pre-clinical imaging modality to non-invasively assess DC-based cancer vaccine efficacy.

## METHODS

### Animal use

C57BL/6 mice were purchased from Charles River Laboratories, Inc. (Senneville, CAN). All animal studies were performed in accordance with institutional and national guidelines.

### Reagents and antibodies

Supplemental Table 1 contains a complete list of reagents and antibodies used in this study.

### Mouse bone marrow-derived dendritic cell (BMDC) generation

BMDC were generated from C57BL/6 mice femurs and tibias as previously detailed [17, 26] with the following modifications. Briefly, BMDC progenitors were cultured in complete RPMI media containing interleukin-(IL)-4 and granulocyte-macrophage colony-stimulating factor (GM-CSF) at 10 ng/mL and 4 ng/mL, respectively for four days at 37ºC and 5% CO_2_. After four days, immature BMDC were enriched from heterogenous cell cultures by Histodenz™ gradient centrifugation.

> *Adenoviral transduction:* Immature day 4 BMDC were transduced with a replication-defective recombinant human adenovirus (Ad) type 5 expressing enhanced green fluorescent protein (eGFP) as a model TAA [27] at a multiplicity of infection (MOI) of 30 for two hours at 37ºC and 5% CO_2_. BMDC incubated in media lacking Ad served as the untransduced control BMDC.
>
> *Superparamagnetic iron oxide (SPIO) labeling and BMDC maturation:* Following Ad transduction, Ad BMDC and untransduced BMDC were labeled as previously described with the SPIO FeREX® (used in Figure 1 only) and Synomag®-D [13, 14, 28]. Briefly, FeREX® alone or Synomag®-D SPIO was complexed with protamine sulfate (0.24 mg/mL) and heparin (8 USP units/mL) in incomplete RPMI and co-incubated with BMDC at a final concentration of 200 µg Fe/mL for four hours at 37ºC and 5% CO_2_. At this point, IL-4-and GM-CSF-supplemented complete RPMI was added to BMDC culture and incubated overnight at 37ºC and 5% CO_2_. On day 5, BMDC were further matured [12] in culture for 24 hours.
>
> FIGURE 1. Ad transduced and SPIO-labeled BMDC are *in vivo* migration competent. CD11c^+^ BMDC were either Ad transduced and then SPIO-labeled (**a**, Ad SPIO^+^, green histogram) or labeled with SPIO and then Ad transduced (**b**, SPIO^+^ Ad, grey histogram). Black histograms in (**a**) and (**b**) depict minimal fluorescence observed with untransduced SPIO^+^ BMDC. Across multiplicities of infection (MOI) of 10, 30 and 60, Ad SPIO^+^ at MOI = 30 resulted in the highest percentage of eGFP^+^ BMDC (**c**, green arrow) and this culture condition was employed for all subsequent experiments. Data is shown as mean ± SEM for n = 3 independent experiments. One million Ad SPIO^+^ BMDC were fluorescently labeled with PKH26 immediately before hind footpad adoptive transfer. Two days later, excised popliteal lymph nodes (pLN) were processed for digital morphometric analysis of PKH26 (**d**, red bar) and eGFP fluorescence (**d**, green bar), with representative fluorescence microscopy images shown in Supplemental Figure 1. The percentage of total BMDC that were eGFP^+^ at the time of injection (**c**, green arrow) is comparable to the percentage of pLN migration competent BMDC that were eGFP^+^ (**d**). Data is representative of n = 2 independent experiments with four mice per group. 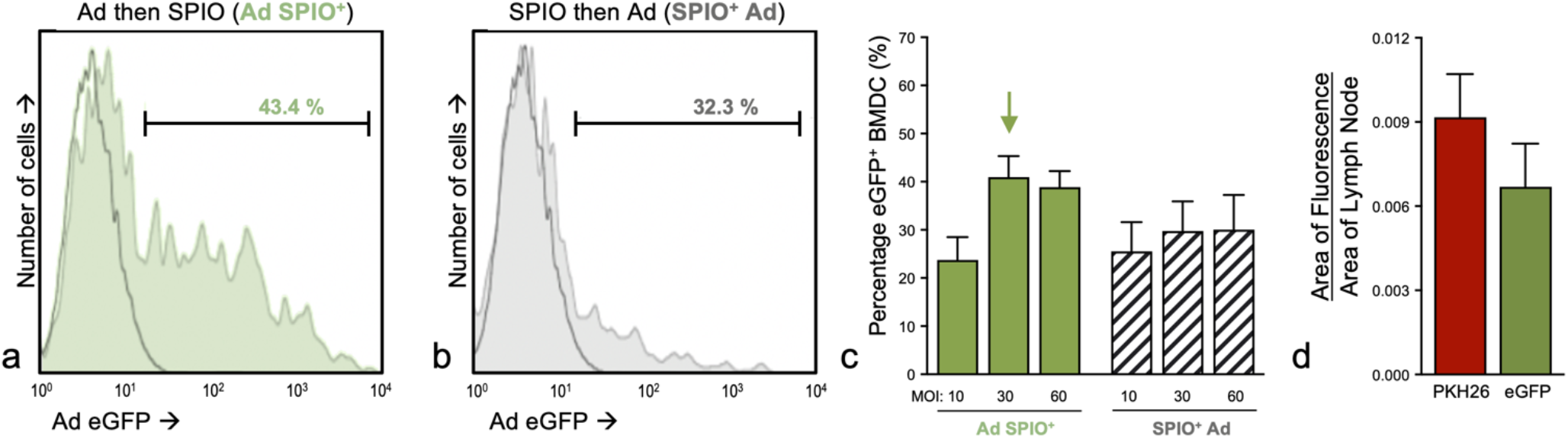
>
> *BMDC magnetic column enrichment:* On the sixth and final culture day, SPIO^+^ BMDC were enriched via magnetic column separation [14]. After collection and repeated washing in PBS to remove free SPIO, BMDC were resuspended in PBS in a 12 × 75 mm polypropylene tube and placed in a EasySep™ magnet (Stemcell Technologies, Vancouver, CAN) for 5 minutes. Both tube and magnet were inverted to collect BMDC not labeled with SPIO (unlabeled, UL) that remained in the flow through. Conversely, both Ad-transduced and untransduced BMDC that incorporated SPIO were retained in the tube and were subsequently collected for downstream application as Ad SPIO^+^ and SPIO^+^ BMDC, respectively.

### Immunophenotyping and viability assessment

A previously used protocol from our group was employed with modifications to assess day 6 mature BMDC viability and phenotype [26] using the fluorescent antibodies and dyes listed in Supplemental Table 1. Zombie NIR™ fixable vital dye was added to BMDC at room temperature for 20 minutes in PBS to determine viability. Next, BMDC underwent surface immunofluorescence staining in the presence of TruStain FcX™ anti-mouse CD16/CD32 block at 4°C for 25 minutes. Extensive washing in HBSS and 0.1% bovine serum albumin ensued prior to fixation in 4% paraformaldehyde and storage at 4°C. When necessary, eGFP expression due to Ad transduction and fluorescent membrane-intercalating dye incorporation (detailed below) was assessed using flow cytometry and compared to an aliquot of non-fluorescent BMDC. Data was acquired on a LSRII analytical flow cytometer (BD Biosciences, San Jose, USA) and analyzed using FlowJo Software (v10, Tree Star, Inc., Ashland, USA).

### Adoptive cell transfer (ACT)

Following magnetic enrichment and immediately prior to ACT, SPIO^+^ and Ad SPIO^+^ BMDC were fluorescently labeled with Tag-It Violet™ (Violet) cell tracking dye (5 µM) using an established protocol for membrane-intercalating dyes [29]. Low (3×10^5^ cell) and high (5×10^5^ cell) injection doses of Violet^+^ SPIO^+^ BMDC were prepared in 40 µL PBS and subcutaneously injected into the right hind footpad of C57BL/6 mice. An identical number of Violet^+^ Ad SPIO^+^ BMDC were injected into the contralateral hind footpad (n=3 mice per injection dose).

In a separate experiment, Ad SPIO^+^ BMDC were labeled with PKH26 as described previously [13] prior to hind footpad ACT of 1×10^6^ PKH26^+^ Ad SPIO^+^ BMDC (n=2 independent experiments with 8 total mice).

### Magnetic particle imaging (MPI)

Mice were imaged as previously described [14] using a Momentum MPI Scanner (Magnetic Insight Inc., Alameda, USA) on days 0 and 2 post injection using imaging parameters described below. Prior to imaging, mice were fasted for 12 hours with water, a laxative, and corn bedding to mitigate gastrointestinal (GI) iron signal in MPI images. Mice were anesthetized initially with 2% isoflurane and maintained with 1% isoflurane during imaging. Afterwards, mice were returned to cages with food *ad libitum*.

> *Day 0:* Post-injection images were acquired in 3D using 3.0 Tesla/m (T/m) selection field gradient and drive field strengths of 20 mT and 23 mT in the X and Z axes, respectively, for isotropic imaging. The MPI signal was collected from 35 projection angles equally distributed between 0° and 180°. A 12 cm (Z) x 6 cm (X) field of view (FOV) was centered on the mouse body and encompassed the entire imaging area without any signal being present at the edges of the FOV. These images were reconstructed using the prescribed native reconstruction algorithm equipped in the Regular User Interface (RUI). The acquisition time for full 12 cm (Z) FOV images was 40 minutes.
>
> *Day 2:* Images were acquired using two different imaging sequences. Initially, mice were imaged using the same 3D image parameters and the reconstruction described above. Subsequently, mice were imaged again in 3D using a 3.0 T/m selection field gradient and drive field strengths of 20 mT and 23 mT in the X and Z axes, respectively, with 35 projection angles. Single channel imaging was used to collect information along the Z axis only. The MPI signal was collected from 35 projection angles equally distributed between 0° and 180°. A smaller 2 cm (Z) x 6 cm (X) FOV was used to only image low signal from BMDC that migrated to pLN and excluded impeding stronger signal from the injection site. These images were reconstructed using a multichannel joint algorithm where an inverse combiner was enabled to prevent image artifacts due to MPI signal being present at the FOV edges [30]. The acquisition time for focused 2 cm (Z) FOV images was 15 minutes. These customized image parameters were implemented using the Advanced User Interface (AUI) which allowed the user access to a configuration editor where changes were made to prescribed sequences equipped on the MPI system. Excised pLN were then imaged using a full 12 cm (Z) FOV.
>
> *Analysis and* quantification: Two samples containing 1×10^6^ SPIO^+^ and 1×10^6^ Ad SPIO^+^ BMDC were singly imaged using the 12 cm (Z) and 2 cm (Z) FOV 3D image parameters described above. Iron loading per cell was measured from these samples by converting the measured MPI signal to iron mass using calibration lines unique to the image sequence, as described [14]. This iron per cell measurement was subsequently used for all *in vivo* and *ex vivo* image analysis to estimate cell numbers from MPI signal. All MPI images were imported into Horos™, an open-source clinically relevant image analysis software (version 3.3.6, Annapolis, USA). Images were viewed using a custom MPI colour look-up table (CLUT). MPI signal was measured within a specific region of interest (ROI) using a semi-automatic segmentation tool. Total MPI signal for an ROI was calculated by multiplying the ROI volume by the mean signal. The signal to noise ratio (SNR) was calculated by dividing the mean signal for a ROI by the standard deviation of the background noise. The SNR had to be greater than 5 for the MPI signal to be considered detectable and for images to be further quantified. Iron content was calculated by dividing the total MPI signal by the slope of the respective calibration line for those image parameters. All MPI images were delineated and analyzed in the same way to ensure consistency. Cell numbers were estimated by dividing the iron quantified within a given ROI by the respective iron/cell measurements.

### Popliteal lymph node (pLN) histology

Upon final *in vivo* MPI imaging session completion, pLN were excised from euthanized mice, fixed with 4% paraformaldehyde, and promptly underwent *ex vivo* MPI. Next, pLN were cryoprotected in sucrose and processed into 16 µm tissue sections [26] for imaging using an EVOS™ M7000 Imaging System (Thermo Fisher Scientific, Burlington, CAN). For quantitative image analysis, pLN sections were imaged for Violet fluorescence. Digital morphometric analysis of imaged pLN sections was performed using Image Pro Premier 9.2 software (Media Cybernetics, Rockville, USA) and represented as area of Violet fluorescence/area of lymph node.

Similarly, to above, pLN were excised and processed from mice that received hind footpad injections of PKH26^+^ Ad SPIO^+^ BMDC two days prior. A Leica DMIRE2 fluorescence microscope (Leica, Wetzlar, GER) and Image Pro software were used to perform digital morphometric analysis of pLN sections [17] to determine the area of PKH26^+^ and eGFP^+^ fluorescence per area of lymph node.

### Statistical analysis

All data is presented as means ± standard error of the mean (SEM). A paired *t*-test, two-way ANOVA with Tukey’s multiple comparison test, and simple linear regression analysis (Graph Pad Prism, Version 9.4.1, La Jolla, USA) were used. Statistical significance was defined when *p* ≤ 0.05.

## RESULTS

### SPIO labeling and BMDC viability are unaffected by prior adenoviral transduction

Although Ad transduction [3] as well as SPIO labeling of dendritic cells [18] are both widely reported on in the context of cancer immunotherapy, the outcome of Ad transduction combined with SPIO labeling within one BMDC generation protocol was unknown. Across a range of Ad MOI, initial experiments demonstrated that BMDC transduction prior to SPIO labeling (Ad^+^ SPIO) consistently transduced 20-45% of BMDC, with decreased transduction percentages observed in BMDC that were SPIO-labeled and then Ad-transduced (SPIO^+^ Ad) (**Fig. 1a-c**). Importantly, the percentage of Ad-transduced BMDC (PKH26^+^ eGFP^+^) of total pLN-migrated Ad SPIO^+^ BMDC (PKH26^+^) was comparable to the percentage of Ad-transduced BMDC at the time of injection (**Fig. 1d & Supplemental Fig. 1**). This indicated that Ad-transduction does not impair *in vivo* BMDC migration. Next, the viability of singlet, CD45^+^CD11c^+^ BMDC was measured using flow cytometry and Zombie NIR™ fixable vital dye (**Fig. 2a-d**). SPIO^+^ and Ad SPIO^+^ BMDC were associated with a small but significant decrease in viability compared to UL BMDC (**Fig. 2e**, grey and green bars compared to white bar, *p* = 0.0008 and *p* = 0.002); however, BMDC viability was consistently >90% across all conditions. More importantly, cell viability was equivalently high for Ad SPIO^+^ BMDC (93.5 ± 0.82%) and for untransduced SPIO^+^ BMDC (93.2 ± 0.74%) (**Fig. 2e**, *p* = 0.823). As expected, UL and untransduced SPIO^+^ BMDC expressed negligible eGFP (**Fig. 2f**, open and grey histogram, respectively) compared to ∼30% of Ad SPIO^+^ BMDC that expressed eGFP (**Fig. 2g**, green histogram). By comparing cell counts prior to and immediately after magnetic column enrichment, the SPIO labeling percentage was calculated and did not significantly differ between SPIO^+^ (78 ± 8.1%) and Ad SPIO^+^ (67 ± 8.6%) BMDC (**Fig. 2h**, *p* = 0.135), suggesting that our currently outlined Ad transduction and SPIO labeling protocol is suitable for continued use.

**FIGURE 2.**
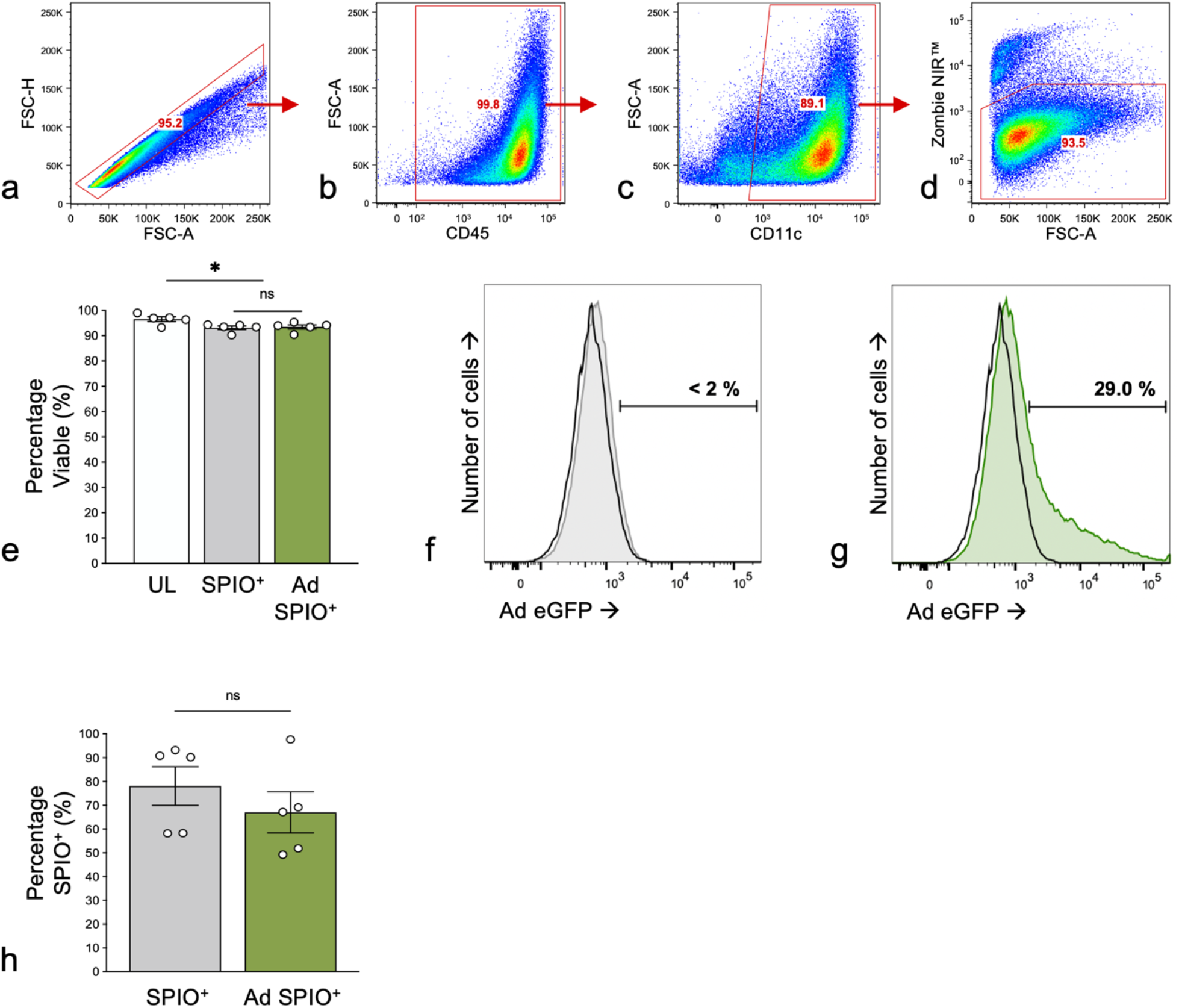
BMDC maintained high viability after Ad transduction and efficiently labeled with SPIO. The viability of singlet (**a**), CD45^+^ (**b**), CD11c^+^ (**c**) BMDC was measured by Zombie NIR™ fixable viability dye and flow cytometry (**d**). No significant differences in viability were measured between SPIO^+^ BMDC (**e**, gray bar) and Ad SPIO^+^ (**e**, green bar), although the viability of both populations exhibited a small but significant decrease compared to untransduced and unlabeled (UL) BMDC (**e**, white bar). The viability was >93% for all conditions. UL (**f**, open histogram) and SPIO^+^ (**f**, gray histogram) BMDC have negligible eGFP expression while Ad SPIO^+^ BMDC express eGFP following transduction with an Ad encoding eGFP (**g**, green histogram). Prior transduction did not affect the percentage of BMDC that successfully labeled with SPIO (**h**). Data are shown as means ± SEM for n = 5 independent experiments (**e**, two-way ANOVA, * *p* = 0.0008 and *p* = 0.002) and means ± SEM for n = 5 independent experiments (**h**, paired *t*-test, ns – no statistical significance, *p* = 0.135).

### CD11c^+^CD86^+^ mature BMDC phenotype unaltered by Ad transduction and SPIO labeling

An important characteristic of a cell labeling agents is efficient incorporation into cells of interest without impacting cell phenotype. Within our set-up, mature BMDC were defined as CD11c^+^CD86^+^ (**Fig. 3a**) and their percent expression did not significantly differ between UL (68.9 ± 4.5%), SPIO^+^ (72.8 ± 2.8%) and Ad SPIO^+^ (74.6 ± 3.2%) BMDC (**Fig. 3b**, white, grey and green bars, respectively). This trend persisted across a range of cell surface markers critical for therapeutic BMDC function and included markers of BMDC migration (**Fig. 3c**, CCR7), activation (**Fig. 3d**, CD40), antigen presentation (**Fig. 3e**, I-A^b^) and T cell stimulation (**Fig. 3f/g**, ICOSL and OX40L). In addition to no phenotypic changes due to Ad transduction or SPIO incorporation, >92% of CD11c^+^CD86^+^ mature BMDC remarkably expressed all phenotypic markers in question, furthering support for our combined Ad transduction and SPIO labeling protocol as we progressed to *in vivo* MPI studies.

**FIGURE 3.**
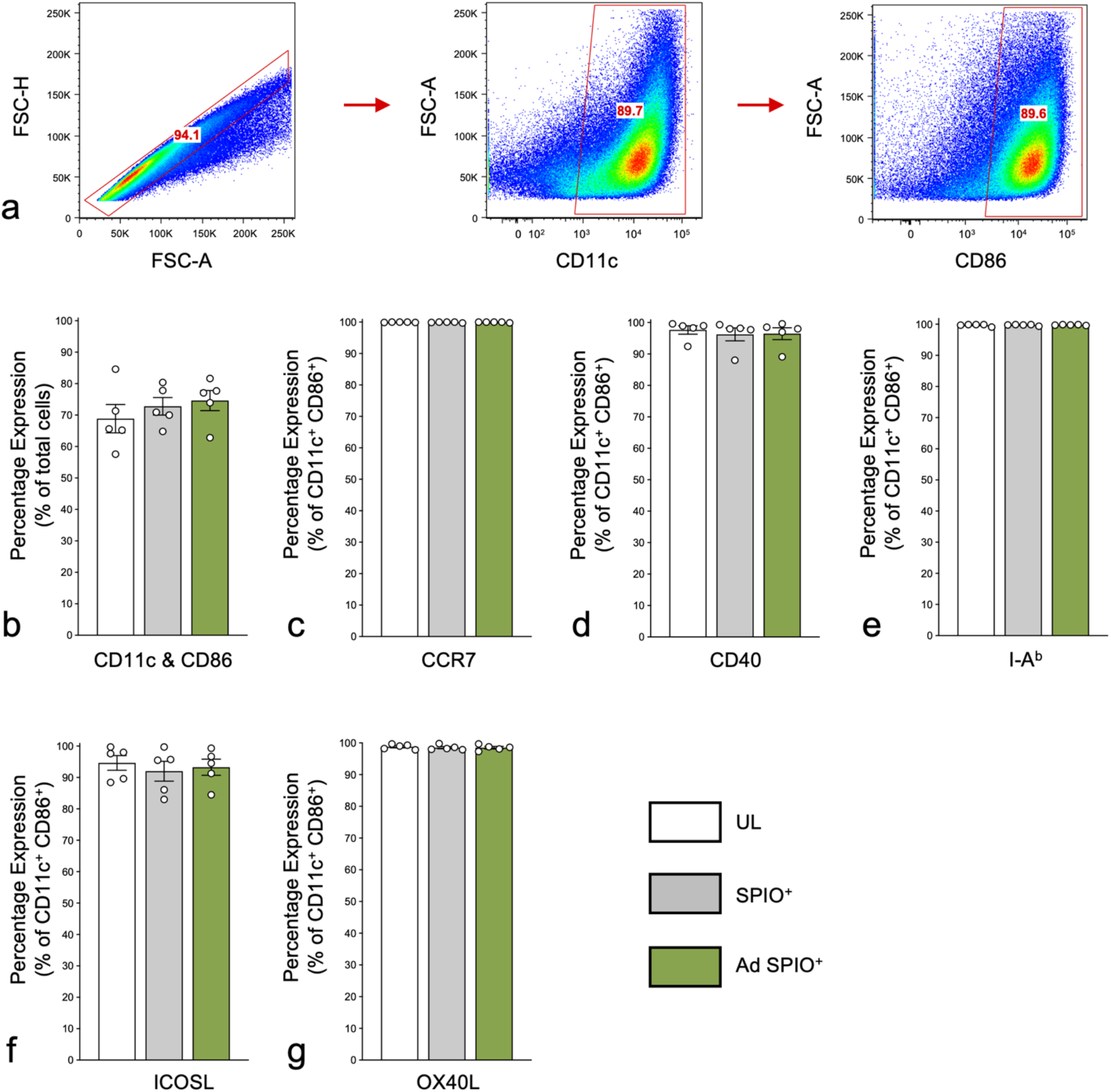
BMDC phenotype is unaffected by SPIO labeling or Ad transduction. Representative flow cytometric dot plots to identify singlet BMDC expressing both CD11c and CD86 (CD11c^+^CD86^+^) are shown in (**a**). The percentage of mature BMDC (identified as CD11c^+^CD86^+^) did not differ between untransduced and unlabeled (UL) BMDC (**b**, white bar), SPIO^+^ BMDC (**b**, gray bar) and Ad SPIO^+^ BMDC (**b**, green bar). Neither SPIO labeling or Ad transduction altered the phenotype of CD11c^+^CD86^+^ mature BMDC as assessed by markers of migration **(c**, CCR7), activation (**d**, CD40), MHC Class II antigen presentation (**e**, I-A^b^) and T cell stimulation (**f** and **g**, ICOSL and OX40L, respectively). Data are shown as means ± SEM for n = 5 independent experiments (**b-g**, two-way ANOVA, * *p* < 0.05).

### *In vivo* detection of SPIO^+^ BMDC and Ad SPIO^+^ BMDC migration using 3D MPI

We next sought to measure SPIO^+^ and Ad SPIO^+^ BMDC *in vivo* migration with MPI. This longitudinal MPI study is outlined schematically in Figure 4 with quantified data that is representative of 5 independent experiments presented in Figure 5.

**FIGURE 4:**
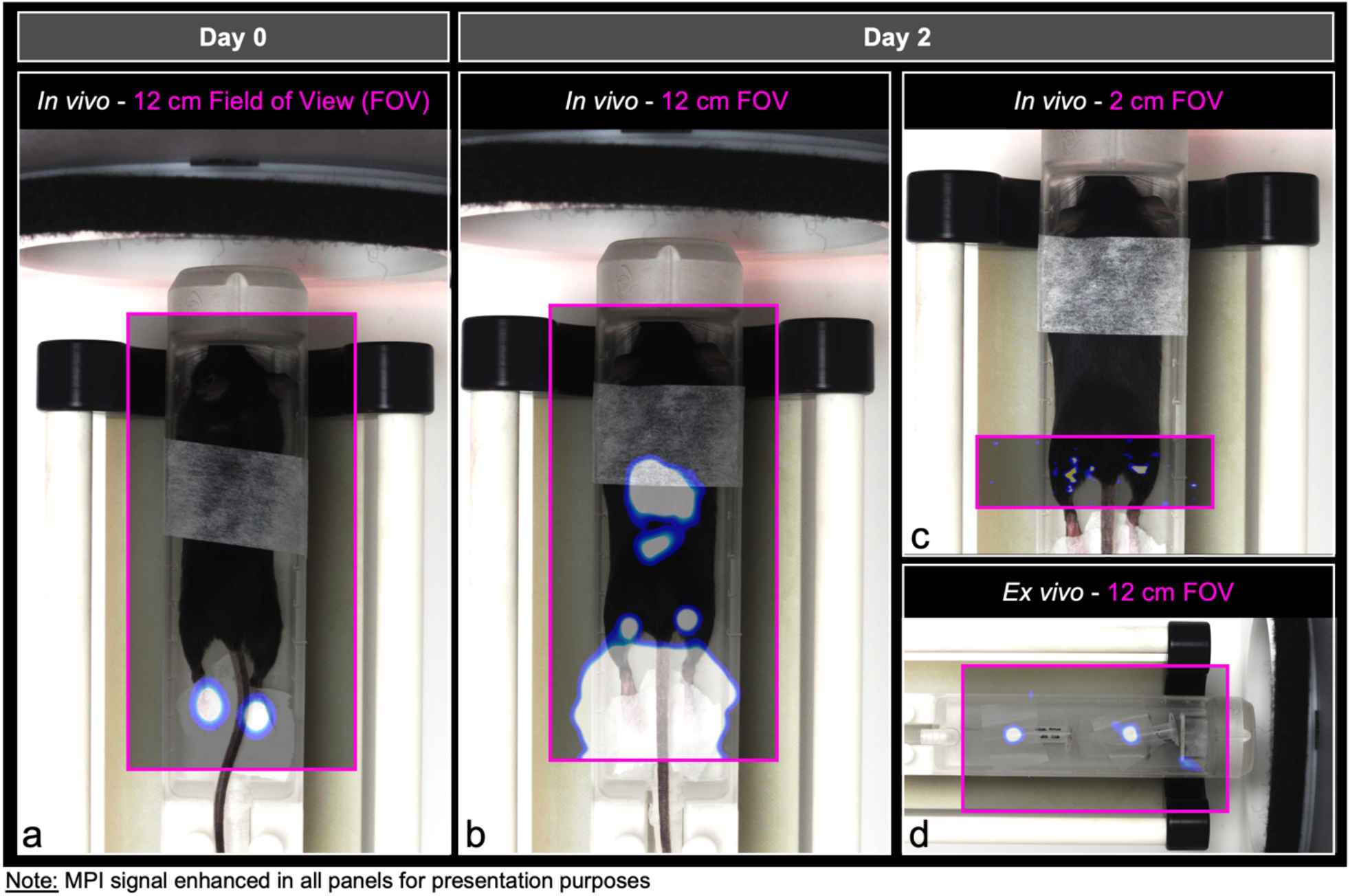
Comparison of FOV size to detect and quantify *in vivo* SPIO^+^ BMDC pLN migration. On day 0, SPIO^+^ and Ad SPIO^+^ BMDC were adoptively transferred into the hind footpads of C57BL/6 mice immediately prior to MPI using a 12 cm whole animal FOV (left panel, pink box). The mouse orientation within the scanner is shown and overlaid with MPI signal. Imaging with a 12 cm FOV (middle panel, pink box) was then repeated two days later and immediately followed by MPI of the pLN region using a 2 cm FOV (upper right panel, pink box). Mice were euthanized to excise pLN that were then placed in a 0.5 mL polypropylene tube before a final *ex vivo* MPI session using a 12 cm FOV (lower right panel, pink box). \*\*\**Note that MPI signal was enhanced in all panels for the purpose of creating a schematic. All other MPI signal presented in this manuscript is not enhanced*.

**FIGURE 5:**
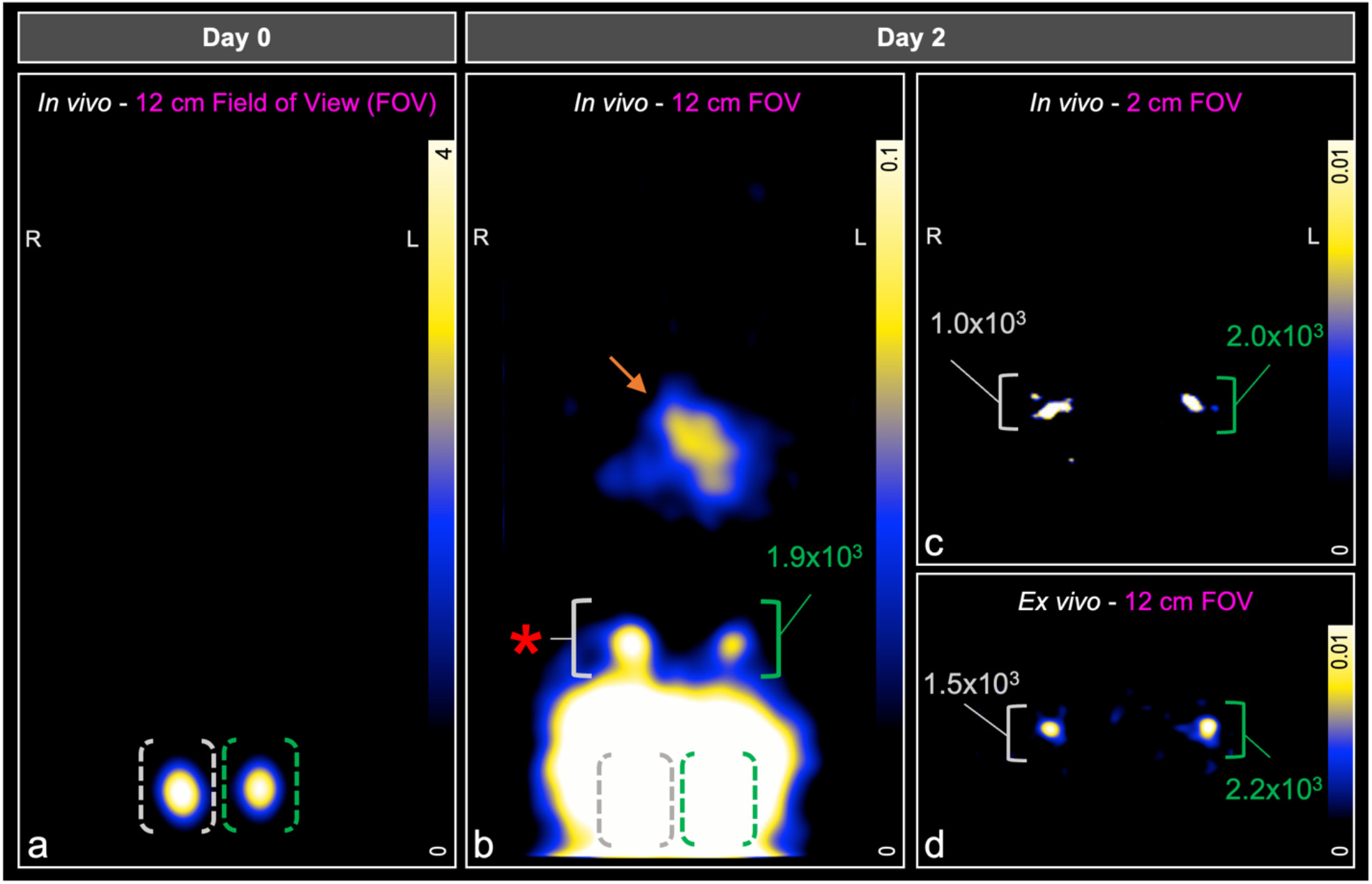
MPI detection and quantification of *in vivo* SPIO^+^ and Ad SPIO^+^ BMDC migration to pLN. On day 0, SPIO^+^ BMDC (**a**, gray outline) and Ad SPIO^+^ BMDC (**a**, green outline) were detected at the injection site (hind footpads, 3×10^5^ BMDC injection dose) immediately following adoptive cell transfer. Two days later, MPI using a 12 cm FOV was unable to quantify SPIO^+^ BMDC migration to the pLN (**b**, gray bracket and red asterisk) but quantified pLN-migrated Ad SPIO^+^ BMDC (**b**, green brackets). GI signal is denoted by the orange arrow in (**b**). In contrast, a 2 cm FOV MPI scan detected and quantified both SPIO^+^ BMDC and Ad SPIO^+^ BMDC migration to the pLN (**c**, gray and green brackets, respectively). pLN were then excised from animals and underwent *ex vivo* MPI to detect pLN-migrated SPIO^+^ BMDC (**d**, gray bracket) and Ad SPIO^+^ BMDC (**d**, green bracket). MPI images are slices selected from 3D imaging using 35 projections. Data is representative of n = 5 independent experiments.

On day 0, scanning was performed immediately after ACT using a 12 cm FOV (**Fig. 4a**) with 3D MPI signal shown in full dynamic range. MPI signal was localized to the right and left hind footpad injection sites due to SPIO^+^ BMDC and Ad SPIO^+^ BMDC, respectively (**Fig. 5a**, white and green bracket, respectively, 5×10^5^ cell injection dose each). Two days later, the same FOV (**Fig. 4b**) and imaging parameters identified a distinct region of interest (ROI) at the left pLN region quantified as 1.9×10^3^ Ad SPIO^+^ (green bracket) but the signal at the right pLN region (white bracket and red asterisk) could not be separated from the high signal emanating from the footpad injection sites (**Fig. 5b**). The 3D MPI signal window leveled to the pLN region results in oversaturation of both GI-localized and persistent injection site signal (**Fig. 5b**, orange arrow, grey and green brackets).

Despite animals being fasted prior to imaging, GI signal was visible due to residual food and iron contained within consumed bedding. To combat the difficulty of resolving pLN signal near a source of higher signal intensity [14], a second day 2 scan using a 2 cm focused FOV (**Fig. 4c**) centred on the pLN was conducted. MPI signal was clearly present in the right and left pLN regions and quantified as 1.0×10^3^ SPIO^+^ and 2.0×10^3^ Ad SPIO^+^ BMDC, respectively (**Fig. 5c**, white and green brackets, respectively). Post-mortem MPI using a 12 cm FOV (**Fig. 4d**) on excised pLN validated *in vivo* signal (**Fig. 5d**, white and green bracket, respectively).

### Focused small FOV MPI with multichannel joint image reconstruction is superior to whole animal FOV MPI for quantifiable detection of pLN-migrated BMDC

For the longitudinal MPI experiment detailed above, day 2 MPI images are shown for all animals that received either a low (**Fig. 6a**, 3×10^5^ cells) or high (**Fig. 6b**, 5×10^5^ cells) injection dose. *In vivo* pLN-localized MPI signal is shown for all animals following acquisition using a 12 cm and focused 2 cm FOV, respectively (**Fig. 6a/b**, left and middle panels, respectively). Excised pLN were also imaged using a 12 cm FOV for *in vivo* validation purposes (**Fig. 6a/b**, right panels). Using a 12 cm FOV, quantifiable signal was detected in only 4/6 right pLN and 5/6 left pLN as summarized for SPIO^+^ and Ad SPIO^+^ BMDC (**Fig. 6c**, grey and green bars). This is in stark contrast to our ability to confidently detect quantifiable signal in 6/6 right pLN and 6/6 left pLN using a focused 2 cm FOV with multichannel joint reconstruction (**Fig. 6d**) and from excised pLN set-up (**Fig. 6e**) that is summarized for SPIO^+^ and Ad SPIO^+^ BMDC (**Fig. 6d/e**, grey and green bars). Irrespective of the FOV employed or the injection dose, there was no significant difference between the number of pLN-migrated SPIO^+^ BMDC and Ad SPIO^+^ BMDC, although the data does trend that increased injection dose leads to increased pLN-migrated BMDC (**Fig. 6c-e**). In summary, we have demonstrated that a focused 2 cm FOV with multichannel joint reconstruction results in quantifiable signal detection in 100% of pLN and is a noted improvement to 75% of pLN with quantifiable signal detected using a 12 cm FOV. It is also important to note that signal quantification using either a 12 cm or 2 cm FOV remains strongly linearly correlated to *ex vivo* pLN quantification (**Fig. 6f/g**, R^2^ = 0.91 and R^2^ = 0.88).

**FIGURE 6:**
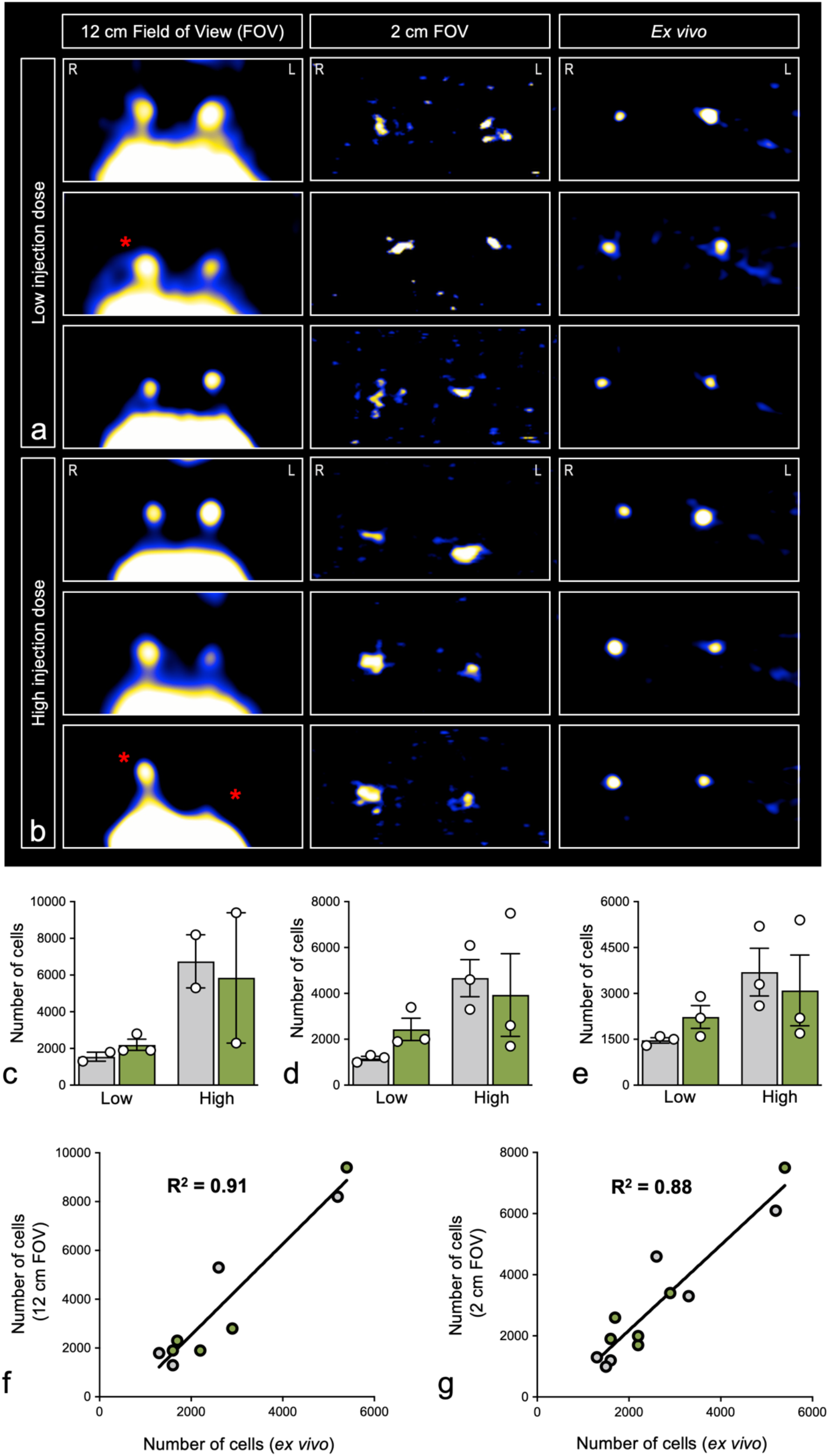
Focused small FOV MPI to accurately quantify *in vivo* BMDC signal. *In vivo* MPI images of the pLN region acquired using a 12 cm FOV (**a**, left panels) and 2 cm FOV (**a**, middle panels) are shown for three mice that received low dose (3×10^5^ SPIO^+^ and Ad SPIO^+^ BMDC) hind footpad injections. Corresponding *ex vivo* pLN MPI images are also shown (**a**, right panels). MPI images of three mice that received high dose (5×10^5^ SPIO^+^ and Ad SPIO^+^ BMDC) injections are displayed in (**b**) using the same layout described in (**a**). Red asterisks indicate pLN in which BMDC migration could not be quantified using a 12 cm FOV (**a** and **b**, left panels) but were quantified using a 2 cm FOV and *ex vivo* MPI set-up (**a** and **b**, middle and right panels, respectively). Images are shown window-leveled to the dynamic range of the pLN. MPI quantification of pLN-migrated BMDC determined using 12 cm FOV (**c**), 2 cm FOV (**d**) and *ex vivo* MPI (**e**) are summarized for low and high dose SPIO^+^ (**c-e**, gray bars) and Ad SPIO^+^ (**c-e**, green bars) BMDC injection conditions. Only signal that was ≥ 5 times the standard deviation above background signal was used for quantification. Linear regression analysis revealed a strong correlation between 12 cm FOV and *ex vivo* pLN MPI quantification (**f**, R^2^ = 0.91) for pLN that were able to be quantified using a 12 cm FOV. In contrast, BMDC migration to all pLN was successfully quantified with a 2 cm FOV and strongly correlated with *ex vivo* pLN MPI quantification (**g**, R^2^ = 0.88). Data is representative of n = 5 independent experiments.

### MPI pLN signal detection is due to originally injected SPIO^+^ and Ad SPIO^+^ BMDC

By rendering both SPIO^+^ and Ad SPIO^+^ BMDC Violet fluorescent (Violet^+^) immediately prior to injection, excised pLN can be imaged after terminal MPI to identify Violet^+^ BMDC and confirm that *in vivo* pLN MPI signal is the result of originally injected BMDC. Compared to an aliquot of BMDC removed prior to labeling, 100% of SPIO^+^ and Ad SPIO^+^ BMDC were Violet^+^. Moreover, the Violet mean fluorescence intensity was consistent across both cell populations and thus, is a suitable readout for comparing pLN migration (**Fig. 7a/b**). Upon completion of day 2 MPI, representative pLN cryosections are shown for Violet^+^ SPIO^+^ and Violet^+^ Ad SPIO^+^ BMDC from both low and high injection doses (**Fig. 7c/e** and **Fig. 7d/f**, respectively). In all instances, Violet^+^ BMDC have localized to the central and paracortical T cell-rich areas of the pLN (**Fig. 7c-f**). Digital morphometry then revealed that SPIO^+^ BMDC and Ad SPIO^+^ BMDC migrate equivalently to the pLN two days post ACT for both low (**Fig. 7g**, p = 0.470) and high (**Fig. 7h**, p = 0.539) injection doses, which is in direct agreement with *in vivo* migration quantified using MPI (**Fig. 6**).

**FIGURE 7.**
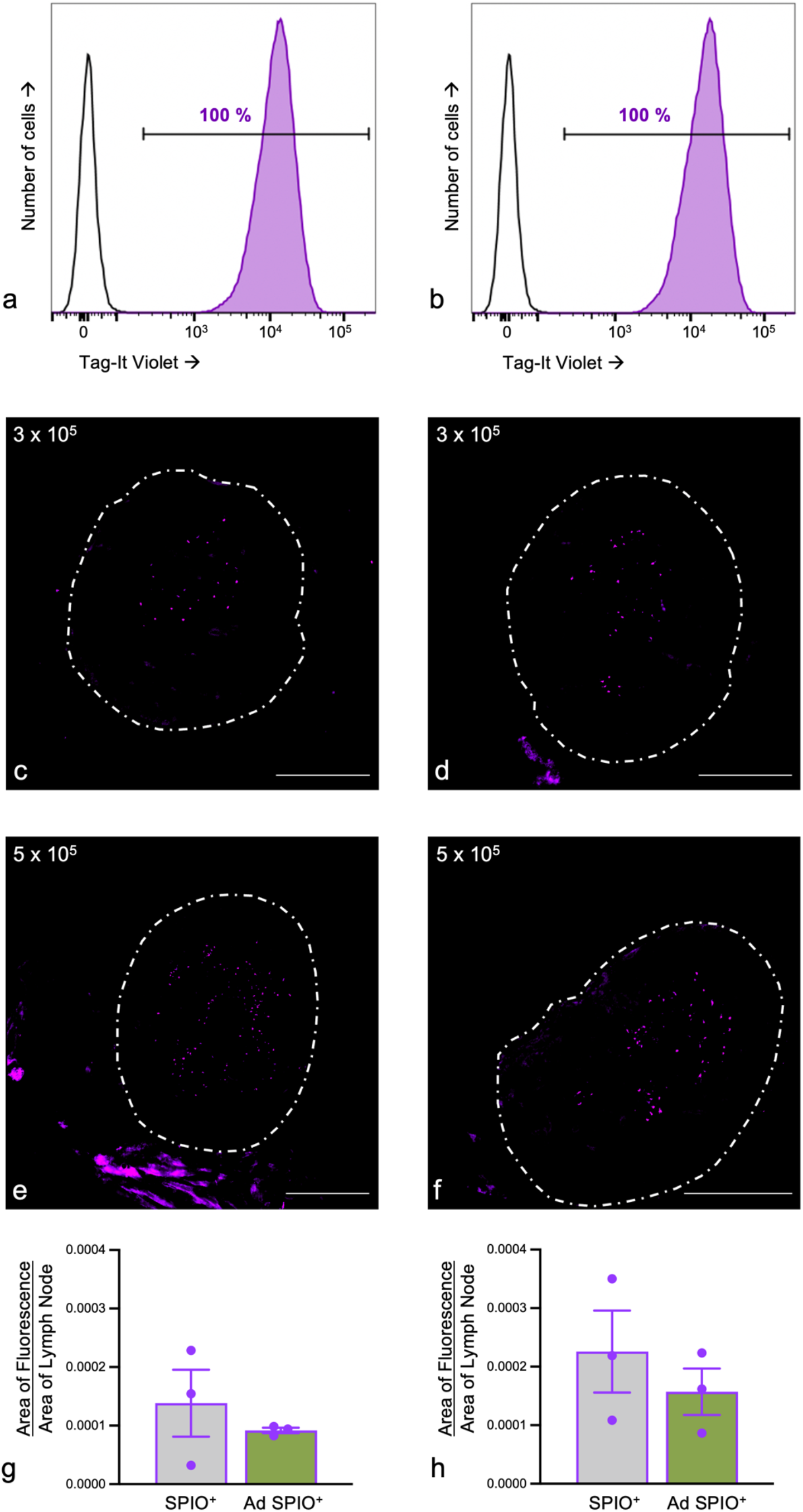
Adoptively transferred SPIO^+^ and Ad SPIO^+^ BMDC are source of MPI signal detected in pLN. Magnetically enriched SPIO^+^ and Ad SPIO^+^ BMDC were labeled with Tag-It Violet (Violet^+^, **a** and **b**, respectively) and formulated into low (3×10^5^) and high (5×10^5^) hind footpad injections. Two days later, pLN were excised and cryosectioned to reveal Violet fluorescence for low (**c**) and high **(e**) SPIO^+^ BMDC and low (**d**) and high (**f**) Ad SPIO^+^ injection doses, respectively. Violet^+^ pLN fluorescence was quantified and did not significantly differ between SPIO^+^ BMDC and Ad SPIO^+^ BMDC for both low (**g**) and high (**h**) injection doses. Data are shown as means ± SEM for n = 3 mice per dose (**g** and **h**, paired *t*-test, *p* = 0.470 and *p* = 0.539). The experimenter was blinded prior to image acquisition and images were taken at 100X magnification (scale bar = 500 μm). Data is representative of n = 5 independent experiments.

## DISCUSSION

For DC-based immunotherapies, the lack of standardization and scientific consensus surrounding administration route, injection dose and optimal DC source [31, 32] means that MPI must be robust enough to detect SPIO-labeled DC regardless of progenitor cell origin. Our prior studies highlighted a limitation of MPI observed when attempting to resolve a strong and weak ROI near each other [14]. To optimize therapeutic DC tracking with MPI, this study describes the efficient labeling of differently generated (Ad-transduced and untransduced) SPIO^+^ BMDC and the use of an image reconstruction algorithm that allows for the choice of a focused FOV to detect and quantify SPIO^+^ DC migration to the pLN 100% of the time despite its nearness to high injection site hind footpad signal.

We have previously reported efficient MRI cell labeling agent uptake by immature DC in the absence of transfection agents [26]; however, transfection agents have been employed to enhance SPIO label uptake and thus, improve MRI/MPI sensitivity to detect therapeutically relevant number of SPIO^+^ BMDC *in vivo* [33]. Despite reports of transfection agents resulting in cytotoxicity [34], protamine sulfate and heparin were successfully used in this study as CD11c^+^ BMDC viability was >90% for SPIO^+^ and Ad SPIO^+^ BMDC (**Fig. 2e**). SPIO nanoparticles for MPI are continuing to be developed with a focus on improving biocompatibility, for example, through surface nanoparticle modifications or liposomal delivery [34, 35] that will improve image resolution and may circumvent the need for transfection agents altogether. Also, we used a naked/dextran-coated SPIO [28] to label DC and is most likely a contributing factor to the reported 70-80% of DC that incorporated SPIO (**Fig. 2h**). Magnetic column enrichment controlled for this reduced labeling percentage by ensuring that only SPIO^+^ and Ad SPIO^+^ DC were characterized and adoptively transferred. Biocompatible SPIO nanoparticles, in addition to improving image resolution [36], will not only improve cell loading but also will improve the percentage of cells that are labeled and is a focus of our future studies.

An important aspect of this study was to demonstrate that BMDC can be dually labeled by Ad and incorporate SPIO nanoparticles without compromising their subsequent ability to migrate *in vivo*. Ad are extensively used for *ex vivo* and *in vivo* DC transduction as they are non-replicating, non-integrating vectors associated with efficient and sustained gene delivery [10, 37] that influences the effectiveness of DC-based cancer vaccines [38]. While eGFP served as a model TAA in this study, Ad transgenes have consisted of a single TAA [39], multiple TAAs to combat antigen loss variants [10] as well as immunostimulatory molecules like CD40, CD137L, 4-1BBL and OX40L [39–42] to activate DC and TAA-specific T cells, respectively. Despite their widespread use, Ad DC transduction still yields conflicting results [43], potentially due to the lack of standardization in generating and the source of therapeutic DC. This may explain why the percentage of Ad-transduced CD11c^+^ BMDC presented here and reported by others [44, 45] is lower than what has been reported in the literature for DC in general [3]. Discrepancies in transduction efficiency between BMDC and DC derived from other sources is primarily attributed to low expression of the coxsackie and adenovirus entry receptors on BMDC [46]. Moreover, we presented a dual Ad transduction SPIO labeling protocol and this combination constrains the modifications that we can make to improve transduction. Ad transduction prior to SPIO labeling was performed based on our earlier studies (**Fig. 1**) and importantly, the presence of heparin during SPIO labeling likely inhibits further transduction after the two-hour initial incubation has elapsed [47]. Extending the transduction incubation time would likely result in increased transduction efficiency but also Ad-induced DC maturation [46] that would lead to maturing DC subsequently incorporating less SPIO.

Although optimizations are necessary (ie. increasing MOI to improve transduction efficiency [43]), this combined protocol is a crucial proof-of-principle study that differently matured DC can efficiently label with SPIO and be tracked *in vivo* with MPI.

Successful DC tracking is contingent on sufficient cell labeling agent incorporation to permit *in vivo* detection without impacting mature DC viability, phenotype, and function [48]. Analysis of BMDC phenotype revealed that neither Ad transduction or SPIO labeling affected CCR7 expression and by extension, BMDC ability to migrate *in vivo*, as nearly 100% of SPIO^+^ mature CD11c^+^CD86^+^ BMDC were CCR7^+^ (**Fig. 3c**). Although additional CCR7-independent markers of DC migration such as CX3CL1 and CXCL12 have been described [49], CCR7 remains the most important DC migration marker. For this reason, prostaglandin E_2_ (PGE_2_) is included in the day 5 maturation cocktail despite its negative effect on IL-12 secretion, which along with IFN-γ, promotes a T_H_1-mediated immune response [32, 50]. In stark contrast to a previous BMDC study [51], we report a remarkably high (near 100%) surface expression of CD40 on SPIO^+^ and Ad SPIO^+^ BMDC (**Fig. 3d**) which should elicit IL-12 production in secondary lymphoid organs following interaction with CD40L on CD4^+^ T_H_ cells [48]. In addition to the CD40:CD40L and I-A^b^:T cell receptor (TCR) interactions between DC and T cells, >92% of BMDC co-express OX40L and ICOSL (**Fig. 3e-g**) to interact with OX40 and ICOS, respectively, on activated T cells [52, 53]. This was expected as TNF-α is a component of the maturation cocktail and induces OX40L and ICOSL upregulation [1, 42]. It is imperative that DC highly express OX40L as interaction with OX40 leads to IFN-γ production [1, 54]. The mature and primed phenotype described above suggests that in a DC-based cancer vaccine setting, both SPIO^+^ and Ad SPIO^+^ BMDC should elicit a strong and sustained T_H_1-mediate response. In conjunction with broad innate immune cell activation conferred by professional APC like DC, this would establish a pro-inflammatory environment conducive for TAA-specific CD8^+^ T cell activation and proliferation [42, 52, 53] that is an absolute requirement for immunotherapeutic effectiveness.

Irrespective of DC source, generation, TAA loading or dose, knowledge of where and in what quantities DC migrate to post injection is required to evaluate DC-based cancer immunotherapy effectiveness [8]. Iron oxide-based MRI has been the most successful non-invasive imaging modality to date for DC tracking owing to the extensive anatomical information it provides alongside its high sensitivity of detection. SPIO^+^ DC *in vivo* produce T2-weighted decreases to SNR capable of detecting as little as ∼2×10^3^ SPIO^+^ cells under pre-clinical ^1^H MRI conditions and permitting comparison of migration differences associated with differently matured DC [12, 18]. Despite its popularity, disadvantages have been widely reported and generally fit into two categories. First, SPIO^+^ DC-induced signal void is not specific and cannot be distinguished from tissues whose chemical shift artifacts generate signal loss [1, 18]. Secondly, as SPIO^+^ DC signal void is not linearly related to iron concentration, it is at best a semi-quantitative imaging modality [17] and is limited in its ability to accurately predict treatment outcome.

Recently, MPI has emerged as an exciting pre-clinical imaging modality capable of directly quantifying the amount of iron, and by extension, the amount of SPIO^+^ cells, within a ROI [14]. This permits for “hot spot” imaging similarly to what is seen for ^19^F MRI [26, 29, 55] and overcomes one of the main limitations of ^1^H MRI detailed above. MPI does not provide any anatomical information; however, it is feasible to combine with a second imaging modality to provide anatomical context to the observed MPI signal [19]. Indeed, a prior study by our group used ^1^H MRI to confirm that SPIO^+^ BMDC detected by MPI were localized to pLN [14] 72 hours following ACT into the hind footpad. This study additionally brought attention to the difficulty of resolving strong and weak MPI signal near each other. We perceive this to be a persistent issue for DC tracking as 3-5% of injected DC reach the pLN (weak signal) [11], which is in close proximity to the hind footpad injection site (strong signal). Even after fasting and housing animals with corn bedding that contains minimal iron prior to MPI, GI signal was still evident (**Figs. 4b&5b**) and could similarly interfere with weak pLN signal quantification. GI signal will also be an issue for human MPI. Therefore, we made use of a focused FOV to quantify pLN signal and exclude areas of high iron concentration that limit the dynamic range capable of being resolved [56].

Two days following ACT, we first employed a standard FOV to measure pLN-migrated SPIO^+^ and Ad SPIO^+^ BMDC. pLN signal was detected in 9 of 12 pLN (75%), which is an improvement to MPI signal detection in 50% of pLN observed in a previous and similarly designed experiment [14]. Strong injection site signal, presumably due to non-migrating or apoptotic BMDC [26], that is unable to be resolved from pLN signal precluded quantification in 3 of 12 pLN. By switching to an advanced image reconstruction algorithm allowing a focused 2 cm FOV that largely excludes injection site MPI signal, pLN signal was detected and quantified in 12 of 12 pLN (100%), demonstrating that MPI is capable of consistent detection and quantification of *in vivo* migration for two distinct DC populations (SPIO^+^ and Ad SPIO^+^ DC). Importantly, a smaller FOV also drastically reduced image acquisition time from 40 minutes to 15 minutes, improving efficiency. We also acknowledge that a focused FOV may not be applicable for imaging studies in which the location of *in vivo* migration is unknown.

Excised pLN MPI was performed in earlier experiments to confirm that MPI signal was originating in the pLN. To remain consistent, we also included excised pLN MPI in this study and as expected, we detected signal in 100% of pLN. Further studies were performed to ensure that pLN-localized signal was due to injected BMDC as opposed to false positive signal resulting from resident macrophage uptake of apoptotic SPIO-labeled cells [57]. By labeling SPIO^+^ and Ad SPIO^+^ BMDC with Violet fluorescence prior to ACT, digital morphometric analysis of pLN confirmed that originally injected migration-competent Violet^+^ DC were the source of MPI signal and that no difference in *in vivo* migration capacity was noted between SPIO^+^ and Ad SPIO^+^ BMDC. In all pLN, SPIO^+^ and Ad SPIO^+^ BMDC penetrated the central and paracortical regions. This suggests that DC were viable upon reaching the pLN as non-viable DC phagocytosed by macrophages would remain in the cortex. Furthermore, this observation has immunological implications as TAA-presenting DC engage with and activate CD4^+^ and CD8^+^ TAA-specific T cells within the paracortex [48, 49] as we demonstrated in a prior BMDC-based cancer vaccine study [26].

Previously, the use of a focused FOV to isolate low signal regions was challenging, if not impossible, with the prescribed native reconstruction algorithm equipped on the Momentum™ MPI scanner and the only option for a user limited to accessing the regular user interface. Native image reconstruction assumes that there is no signal along the edges of the FOV; if there is signal present, these values are set to zero for each line along the transmit axis. This assumption causes an inverted negative image artifact which can prevent signal detection and quantification. Thus, in previous experiments, a full 12 cm FOV was used to ensure all signal was encompassed within a large area. For these experiments, our MPI system was upgraded to allow the user access to an advanced user interface where image sequences could be changed through the configuration editor. Most importantly, this permitted the use of a novel reconstruction algorithm referred to as multichannel joint image reconstruction [30], recently implemented as a reconstruction option on the Momentum™ scanner. The multichannel joint reconstruction method uses an iterative reconstruction technique to recover edge information using information from an orthogonal axis, preventing this artifact, and allowing the user to prescribe a small FOV on the region of interest. Additionally, with this improved method, as few as 1000 pLN-migrated DC (4.4 ng Fe) were detected *in vivo* following footpad injection. This is the lowest amount of Fe detected *in vivo* thus far and the lowest number of cells detected that were labeled with a commercially available SPIO. Previously, Song *et al.* reported on the *in vivo* detection of 250 HeLa cells (7.8 ng Fe) labeled with custom synthesized Janus particles and implanted subcutaneously on the back of a mouse [60]. This study demonstrates the feasibility of using this type of reconstruction to resolve and quantify discrete sources of signal without interference from proximate, varying iron concentrations which are not of interest.

## Conclusions

This is the first *in vivo* MPI study that employed multichannel joint image reconstruction with a focused FOV for image acquisition and represents an important step forward in establishing MPI as a non-invasive imaging modality capable of tracking the fate of injected DC to serve as a surrogate marker of therapeutic effectiveness. Recently, DC-based therapies have garnered much interest within combination cancer immunotherapy in autoimmune and transplantation settings [58] as well as in infectious disease settings such as SARS-CoV-2 [59]. Regardless of disease setting, non-invasive and quantifiable imaging is required for cell-based immunotherapies to be successful.

## Acknowledgments

The authors acknowledge Patrick Goodwill and Justin Konkle for providing the reconstruction technique and assisting with its implementation on the Momentum™ scanner.

## REFERENCES

1. Gao Y, Wang Z, Cui Y, Xu M, Weng L (2022) Emerging Strategies of Engineering and Tracking Dendritic Cells for Cancer Immunotherapy. ACS Appl Bio Mater. https://doi.org/10.1021/ACSABM.2C00790

2. Datta J, Terhune JH, Lowenfeld L, et al (2014) Optimizing Dendritic Cell-Based Approaches for Cancer Immunotherapy. Yale J Biol Med 87:491

3. Lee CS, Bishop ES, Zhang R, et al (2017) Adenovirus-Mediated Gene Delivery: Potential Applications for Gene and Cell-Based Therapies in the New Era of Personalized Medicine. Genes Dis 4:43–63. https://doi.org/10.1016/J.GENDIS.2017.04.001

4. Waldmann TA (2018) Cytokines in Cancer Immunotherapy. Cold Spring Harb Perspect Biol 10:. https://doi.org/10.1101/CSHPERSPECT.A028472

5. Anguille S, Smits EL, Lion E, van Tendeloo VF, Berneman ZN (2014) Clinical use of dendritic cells for cancer therapy. Lancet Oncol 15:. https://doi.org/10.1016/S1470-2045(13)70585-0

6. Butterfield LH (2013) Dendritic cells in cancer immunotherapy clinical trials: are we making progress? Front Immunol 4:. https://doi.org/10.3389/FIMMU.2013.00454

7. Förster R, Braun A, Worbs T (2012) Lymph node homing of T cells and dendritic cells via afferent lymphatics. Trends Immunol 33:271–280. https://doi.org/10.1016/J.IT.2012.02.007

8. Martín-Fontecha A, Sebastiani S, Höpken UE, et al (2003) Regulation of dendritic cell migration to the draining lymph node: impact on T lymphocyte traffic and priming. J Exp Med 198:615–621. https://doi.org/10.1084/JEM.20030448

9. Scheid E, Major P, Bergeron A, et al (2016) Tn-MUC1 DC Vaccination of Rhesus Macaques and a Phase I/II Trial in Patients with Nonmetastatic Castrate-Resistant Prostate Cancer. Cancer Immunol Res 4:881–892. https://doi.org/10.1158/2326-6066.CIR-15-0189

10. Blalock LAT, Landsberg J, Messmer MN, et al (2012) Human dendritic cells adenovirally-engineered to express three defined tumor antigens promote broad adaptive and innate immunity. Oncoimmunology 1:287–357. https://doi.org/10.4161/ONCI.18628

11. Dekaban GA, Hamilton AM, Fink CA, et al (2013) Tracking and evaluation of dendritic cell migration by cellular magnetic resonance imaging. Wiley Interdiscip Rev Nanomed Nanobiotechnol 5:469–483. https://doi.org/10.1002/WNAN.1227

12. de Chickera S, Willert C, Mallet C, Foley R, Foster P, Dekaban GA (2012) Cellular MRI as a suitable, sensitive non-invasive modality for correlating in vivo migratory efficiencies of different dendritic cell populations with subsequent immunological outcomes. Int Immunol 24:29–41. https://doi.org/10.1093/INTIMM/DXR095

13. de Chickera SN, Snir J, Willert C, et al (2011) Labelling dendritic cells with SPIO has implications for their subsequent in vivo migration as assessed with cellular MRI. Contrast Media Mol Imaging 6:314–327. https://doi.org/10.1002/CMMI.433

14. Gevaert JJ, Fink C, Dikeakos JD, Dekaban GA, Foster PJ (2022) Magnetic Particle Imaging Is a Sensitive In Vivo Imaging Modality for the Detection of Dendritic Cell Migration. Mol Imaging Biol 24:. https://doi.org/10.1007/S11307-022-01738-W

15. de Vries IJM, Lesterhuis WJ, Barentsz JO, et al (2005) Magnetic resonance tracking of dendritic cells in melanoma patients for monitoring of cellular therapy. Nat Biotechnol 23:1407–1413. https://doi.org/10.1038/NBT1154

16. Heyn C, Ronald JA, Mackenzie LT, et al (2006) In vivo magnetic resonance imaging of single cells in mouse brain with optical validation. Magn Reson Med 55:23–29. https://doi.org/10.1002/MRM.20747

17. Dekaban GA, Snir J, Shrum B, et al (2009) Semiquantitation of mouse dendritic cell migration in vivo using cellular MRI. J Immunother 32:240–251. https://doi.org/10.1097/CJI.0B013E318197B2A0

18. Bulte JWM, Shakeri-Zadeh A (2022) In Vivo MRI Tracking of Tumor Vaccination and Antigen Presentation by Dendritic Cells. Mol Imaging Biol 24:198–207. https://doi.org/10.1007/S11307-021-01647-4

19. Zheng B, von See MP, Yu E, et al (2016) Quantitative Magnetic Particle Imaging Monitors the Transplantation, Biodistribution, and Clearance of Stem Cells In Vivo. Theranostics 6:291–301. https://doi.org/10.7150/THNO.13728

20. Paysen H, Loewa N, Stach A, et al (2020) Cellular uptake of magnetic nanoparticles imaged and quantified by magnetic particle imaging. Sci Rep 10:. https://doi.org/10.1038/S41598-020-58853-3

21. Graeser M, Knopp T, Szwargulski P, et al (2017) Towards Picogram Detection of Superparamagnetic Iron-Oxide Particles Using a Gradiometric Receive Coil. Sci Rep 7:. https://doi.org/10.1038/S41598-017-06992-5

22. Poller WC, Löwa N, Wiekhorst F, et al (2016) Magnetic Particle Spectroscopy Reveals Dynamic Changes in the Magnetic Behavior of Very Small Superparamagnetic Iron Oxide Nanoparticles During Cellular Uptake and Enables Determination of Cell-Labeling Efficacy. J Biomed Nanotechnol 12:337–346. https://doi.org/10.1166/JBN.2016.2204

23. Knopp T, Gdaniec N, Möddel M (2017) Magnetic particle imaging: from proof of principle to preclinical applications. Phys Med Biol 62:R124–R178. https://doi.org/10.1088/1361-6560/AA6C99

24. Liu S, Chiu-Lam A, Rivera-Rodriguez A, et al (2021) Long circulating tracer tailored for magnetic particle imaging. Nanotheranostics 5:348–361. https://doi.org/10.7150/NTNO.58548

25. Graeser M, Thieben F, Szwargulski P, et al (2019) Human-sized magnetic particle imaging for brain applications. Nat Commun 10:. https://doi.org/10.1038/S41467-019-09704-X

26. Fink C, Smith M, Gaudet JM, Makela A, Foster PJ, Dekaban GA (2020) Fluorine-19 Cellular MRI Detection of In Vivo Dendritic Cell Migration and Subsequent Induction of Tumor Antigen-Specific Immunotherapeutic Response. Mol Imaging Biol 22:549–561. https://doi.org/10.1007/S11307-019-01393-8

27. Damjanovic D, Zhang X, Mu J, Fe Medina M, Xing Z (2008) Organ distribution of transgene expression following intranasal mucosal delivery of recombinant replication-defective adenovirus gene transfer vector. Genet Vaccines Ther 6:. https://doi.org/10.1186/1479-0556-6-5

28. Vogel P, Kampf T, Rückert MA, et al (2021) Synomag®: The new high-performance tracer for magnetic particle imaging. Int J Magn Part Imaging 7:. https://doi.org/10.18416/IJMPI.2021.2103003

29. Fink C, Gaudet JM, Fox MS, et al (2018) 19F-perfluorocarbon-labeled human peripheral blood mononuclear cells can be detected in vivo using clinical MRI parameters in a therapeutic cell setting. Sci Rep 8:. https://doi.org/10.1038/S41598-017-19031-0

30. Konkle JJ, Goodwill PW, Hensley DW, Orendorff RD, Lustig M, Conolly SM (2015) A Convex Formulation for Magnetic Particle Imaging X-Space Reconstruction. PLoS One 10:. https://doi.org/10.1371/JOURNAL.PONE.0140137

31. Santos PM, Butterfield LH (2018) Dendritic Cell-Based Cancer Vaccines. J Immunol 200:71–74. https://doi.org/10.4049/JIMMUNOL.1701024

32. Nava S, Lisini D, Frigerio S, Bersano A (2021) Dendritic Cells and Cancer Immunotherapy: The Adjuvant Effect. Int J Mol Sci 22:. https://doi.org/10.3390/IJMS222212339

33. Kobukai S, Baheza R, Cobb JG, et al (2010) Magnetic nanoparticles for imaging dendritic cells. Magn Reson Med 63:1383–1390. https://doi.org/10.1002/MRM.22313

34. Nejadnik H, Jung KO, Theruvath AJ, et al (2020) Instant labeling of therapeutic cells for multimodality imaging. Theranostics 10:6024–6034. https://doi.org/10.7150/THNO.39554

35. Dadfar SM, Roemhild K, Drude NI, et al (2019) Iron oxide nanoparticles: Diagnostic, therapeutic and theranostic applications. Adv Drug Deliv Rev 138:302–325. https://doi.org/10.1016/J.ADDR.2019.01.005

36. Yang X, Shao G, Zhang Y, et al (2022) Applications of Magnetic Particle Imaging in Biomedicine: Advancements and Prospects. Front Physiol 13:. https://doi.org/10.3389/FPHYS.2022.898426

37. Mendell JR, Al-Zaidy SA, Rodino-Klapac LR, et al (2021) Current Clinical Applications of In Vivo Gene Therapy with AAVs. Mol Ther 29:464–488. https://doi.org/10.1016/J.YMTHE.2020.12.007

38. Montico B, Lapenta C, Ravo M, et al (2017) Exploiting a new strategy to induce immunogenic cell death to improve dendritic cell-based vaccines for lymphoma immunotherapy. Oncoimmunology 6:. https://doi.org/10.1080/2162402X.2017.1356964

39. Hangalapura BN, Oosterhoff D, Aggarwal S, et al (2010) Selective transduction of dendritic cells in human lymph nodes and superior induction of high-avidity melanoma-reactive cytotoxic T cells by a CD40-targeted adenovirus. J Immunother 33:706–715. https://doi.org/10.1097/CJI.0B013E3181ECCBD4

40. Ding J, Jiang N, Zheng Y, et al (2022) Adenovirus vaccine therapy with CD137L promotes CD8+ DCs-mediated multifunctional CD8+ T cell immunity and elicits potent anti-tumor activity. Pharmacol Res 175:. https://doi.org/10.1016/J.PHRS.2021.106034

41. Wenthe J, Naseri S, Hellström AC, et al (2022) Immune priming using DC- and T cell-targeting gene therapy sensitizes both treated and distant B16 tumors to checkpoint inhibition. Mol Ther Oncolytics 24:429–442. https://doi.org/10.1016/J.OMTO.2022.01.003

42. Ylösmäki E, Ylösmäki L, Fusciello M, et al (2021) Characterization of a novel OX40 ligand and CD40 ligand-expressing oncolytic adenovirus used in the PeptiCRAd cancer vaccine platform. Mol Ther Oncolytics 20:459–469. https://doi.org/10.1016/J.OMTO.2021.02.006

43. Sun QF, Zhao XN, Peng CL, et al (2015) Immunotherapy for Lewis lung carcinoma utilizing dendritic cells infected with CK19 gene recombinant adenoviral vectors. Oncol Rep 34:2289–2295. https://doi.org/10.3892/OR.2015.4231

44. Tatsis N, Ertl HCJ (2004) Adenoviruses as vaccine vectors. Mol Ther 10:616–629. https://doi.org/10.1016/J.YMTHE.2004.07.013

45. Mercier S, Gahéry-Segard H, Monteil M, et al (2002) Distinct roles of adenovirus vector-transduced dendritic cells, myoblasts, and endothelial cells in mediating an immune response against a transgene product. J Virol 76:2899–2911. https://doi.org/10.1128/JVI.76.6.2899-2911.2002

46. Strack A, Deinzer A, Thirion C, et al (2022) Breaking Entry-and Species Barriers: LentiBOOST® Plus Polybrene Enhances Transduction Efficacy of Dendritic Cells and Monocytes by Adenovirus 5. Viruses 14:. https://doi.org/10.3390/V14010092

47. Cheng C, Gall JGD, Kong WP, et al (2007) Mechanism of ad5 vaccine immunity and toxicity: fiber shaft targeting of dendritic cells. PLoS Pathog 3:0239–0245. https://doi.org/10.1371/JOURNAL.PPAT.0030025

48. Wimmers F, Schreibelt G, Sköld AE, Figdor CG, de Vries IJM (2014) Paradigm Shift in Dendritic Cell-Based Immunotherapy: From in vitro Generated Monocyte-Derived DCs to Naturally Circulating DC Subsets. Front Immunol 5:. https://doi.org/10.3389/FIMMU.2014.00165

49. Feng M, Zhou S, Yu Y, Su Q, Li X, Lin W (2021) Regulation of the Migration of Distinct Dendritic Cell Subsets. Front Cell Dev Biol 9:. https://doi.org/10.3389/FCELL.2021.635221

50. Jonuleit H, Kühn U, Müller G, et al (1997) Pro-inflammatory cytokines and prostaglandins induce maturation of potent immunostimulatory dendritic cells under fetal calf serum-free conditions. Eur J Immunol 27:3135–3142. https://doi.org/10.1002/EJI.1830271209

51. Zhang C, Xu Z, Di H, Zeng E, Jiang Y, Liu D (2020) Gadolinium-doped Au@prussian blue nanoparticles as MR/SERS bimodal agents for dendritic cell activating and tracking. Theranostics 10:6061–6071. https://doi.org/10.7150/THNO.42114

52. Simonetta F, Alam IS, Lohmeyer JK, et al (2021) Molecular Imaging of Chimeric Antigen Receptor T Cells by ICOS-ImmunoPET. Clin Cancer Res 27:1058–1068. https://doi.org/10.1158/1078-0432.CCR-20-2770

53. Liu C, Zhang Z, Ping Y, et al (2020) Comprehensive Analysis of PD-1 Gene Expression, Immune Characteristics and Prognostic Significance in 1396 Glioma Patients. Cancer Manag Res 12:4399–4410. https://doi.org/10.2147/CMAR.S238174

54. Liu C, Lou Y, Lizée G, et al (2008) Plasmacytoid dendritic cells induce NK cell-dependent, tumor antigen-specific T cell cross-priming and tumor regression in mice. J Clin Invest 118:1165–1175. https://doi.org/10.1172/JCI33583

55. Fink C, Smith M, Sehl OC, et al (2020) Quantification and characterization of granulocyte macrophage colony-stimulating factor activated human peripheral blood mononuclear cells by fluorine-19 cellular MRI in an immunocompromised mouse model. Diagn Interv Imaging 101:577–588. https://doi.org/10.1016/J.DIII.2020.02.004

56. Boberg M, Gdaniec N, Szwargulski P, Werner F, Möddel M, Knopp T (2021) Simultaneous imaging of widely differing particle concentrations in MPI: problem statement and algorithmic proposal for improvement. Phys Med Biol 66:. https://doi.org/10.1088/1361-6560/ABF202

57. Gonzales C, Yoshihara HAI, Dilek N, et al (2016) In-Vivo Detection and Tracking of T Cells in Various Organs in a Melanoma Tumor Model by 19F-Fluorine MRS/MRI. PLoS One 11:. https://doi.org/10.1371/JOURNAL.PONE.0164557

58. Cooke F, Neal M, Wood MJ, et al (2022) Fluorine labelling of therapeutic human tolerogenic dendritic cells for 19F-magnetic resonance imaging. Front Immunol 13: https://doi.org/10.3389/FIMMU.2022.988667

59. Pérez-Gómez A, Vitallé J, Gasca-Capote C, et al (2021) Dendritic cell deficiencies persist seven months after SARS-CoV-2 infection. Cell Mol Immunol 18:2128–2139. https://doi.org/10.1038/S41423-021-00728-2

60. Song G, Chen M, Zhang Y, et al (2018) Janus Iron Oxides @ Semiconducting Polymer Nanoparticle Tracer for Cell Tracking by Magnetic Particle Imaging. Nano Lett. 18, 182–18. https://doi.org/10.1021/acs.nanolett.7b03829

