## Supplemental Table 1 for "*In vivo* tracking of adenoviral-transduced iron oxide-labeled bone marrow-derived dendritic cells using magnetic particle imaging"

**Supplemental Table 1.** Antibodies and reagents.

| Target | Fluorochrome | Company | Clone |
|---|---|---|---|
| B220 (purified) | N/A | BD Biosciences (Mississauga, CAN) | RA3-6B2 |
| CCR7 | PE | Biolegend (San Diego, USA) | 4B12 |
| CD11c | APC | Biolegend | N418 |
| CD40 | PE-Cy5 | Biolegend | 3/23 |
| CD45 | PE | Biolegend | 30-F11 |
| CD86 | PerCP | Biolegend | GL-1 |
| I-Ab | APC-Fire™ 750 | Biolegend | AF6-120.1 |
| I-Ab (purified) | N/A | Biolegend | 25-9-17 |
| ICOS Ligand | PE | Biolegend | HK5.3 |
| OX40 Ligand | PE-Cy7 | Biolegend | RM134L |
| TruStain FcX™ | N/A | Biolegend | 93 |
| Cell proliferation dye | Tag-It Violet™ | Biolegend | N/A |
| Fixable vital dye | Zombie NIR™ | Biolegend | N/A |

| Reagent | Company |
|---|---|
| 2-Mercaptoethanol | Thermo Fisher Scientific (Burlington, CAN) |
| Ad eGFP | Vector Biolabs (Malvern, USA) |
| BSA | Millipore Sigma (Burlington, USA) |
| CpG ODN 1826 | InvivoGen (San Diego, USA) |
| FBS | Thermo Fisher Scientific |
| FeREX® | Biopal Inc. (Worcester, MA, USA) |
| GM-CSF (recombinant) | PeproTech (Montreal, CAN) |
| HBSS | Thermo Fisher Scientific |
| Heparin sodium injection USP | Sandoz Canada Inc. (Boucherville, CAN) |
| HEPES (1 M) | Thermo Fisher Scientific |
| Histodenz™ | Millipore Sigma |
| Interleukin- (IL)-1β | PeproTech |
| IL-4 (recombinant) | PeproTech |
| IL-6 (recombinant) | PeproTech |
| MEM non-essential amino acids solution (100X) | Thermo Fisher Scientific |
| PBS | Thermo Fisher Scientific |
| Penicillin-Streptomycin-L-Glutamine (100X) | Thermo Fisher Scientific |
| PKH26 red fluorescent cell membrane label | Millipore Sigma |
| Prostaglandin E2 | Millipore Sigma |
| Protamine sulfate | Millipore Sigma |
| Rabbit complement (standard) | Cedarlane Labs (Burlington, CAN) |

| | |
|---|---|
| RPMI media | Thermo Fisher Scientific |
| Sodium pyruvate (100 mM) | Thermo Fisher Scientific |
| Synomag®-D | Micromod GmbH (Rostock, GER) |
| TNF-α (recombinant) | PeproTech |

**Ad**, adenovirus; **eGFP**, enhanced green fluorescent protein; **BSA**, bovine serum albumin; **FBS**, fetal bovine serum; **GM-CSF**, granulocyte-macrophage colony-stimulating factor; **HBSS**, Hank's balanced salt solution; **HEPES**, 4-(2-hydroxyethyl)-1-piperazineethanesulfonic acid; **MEM**, minimal essential media; **PBS**, phosphate-buffered saline; **RPMI media**, Roswell Park Memorial Institute media; **TNF**, tumor necrosis factor.
