## Supplemental Figure 1 for "*In vivo* tracking of adenoviral-transduced iron oxide-labeled bone marrow-derived dendritic cells using magnetic particle imaging"

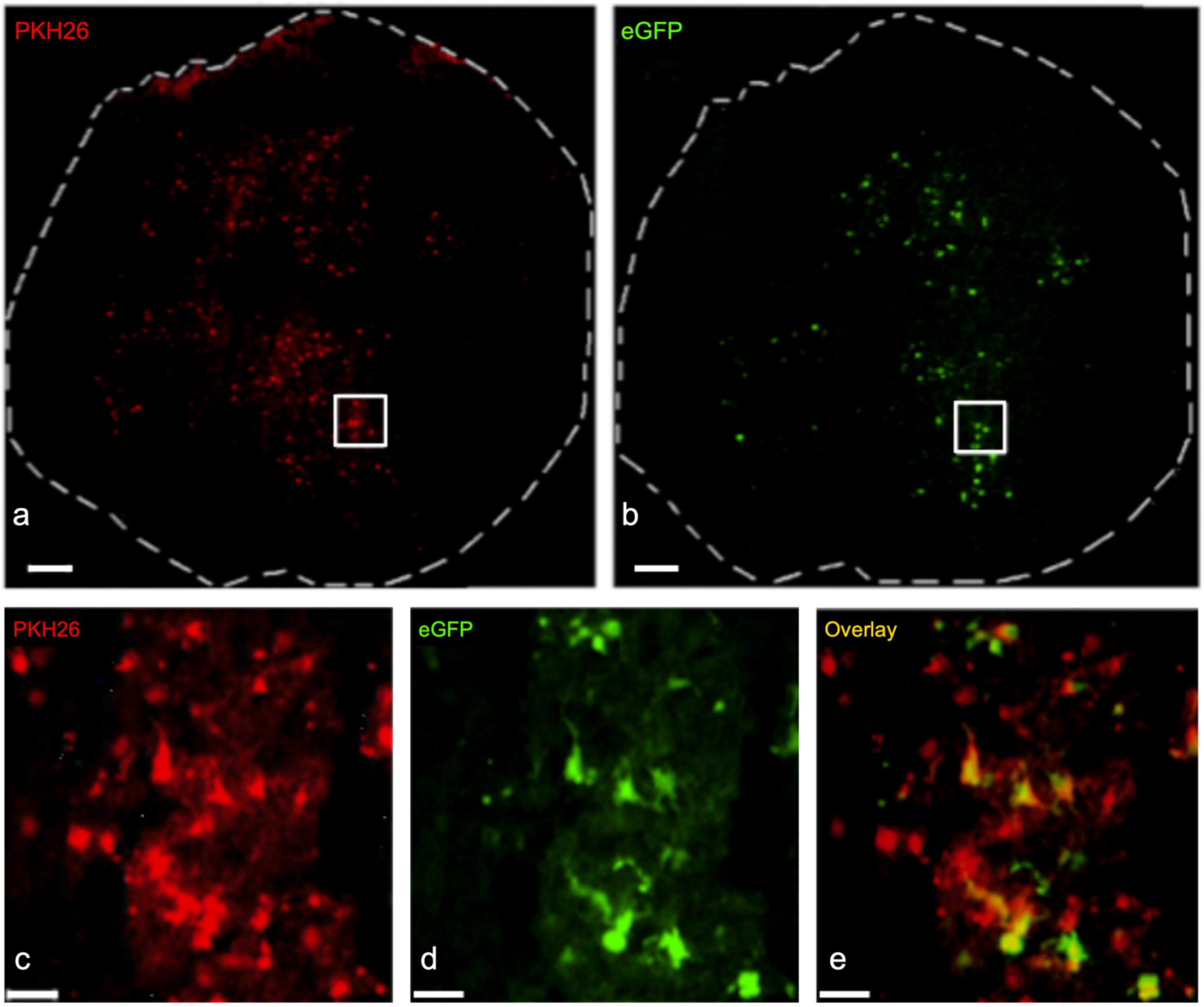


**SUPPLEMENTAL FIGURE 1. Ad SPIO$^+$ BMDC migrate to the popliteal lymph node following injection.** One million Ad SPIO$^+$ BMDC were fluorescently labeled with PKH26 immediately before hind footpad adoptive transfer. Two days later, excised popliteal lymph nodes (pLN) were processed into 16 µm cryosections to observe PKH26 (**a**) and eGFP fluorescence (**b**) at 10X magnification (scale bars = 100 µm). Image inset outlined by a white square in (**a&b**) is shown at 40X magnification for PKH26 (**c**) and eGFP fluorescence (**d**) as well as a PKH26 and eGFP overlay (**e**, yellow) to identify Ad SPIO$^+$ BMDC in pLN. Scale bars = 20 µm in (**c-e**). Data is representative of n=2 independent experiments with four mice per group.
